# SPCS: A Spatial and Pattern Combined Smoothing Method for Spatial Transcriptomic Expression

**DOI:** 10.1101/2021.11.02.467030

**Authors:** Yusong Liu, Tongxin Wang, Ben Duggan, Michael Sharpnack, Kun Huang, Jie Zhang, Xiufen Ye, Travis S. Johnson

## Abstract

High dimensional, localized RNA sequencing is now possible owing to recent developments in spatial transcriptomics (ST). ST is based on highly multiplexed sequence analysis and uses barcodes to match the sequenced reads to their respective tissue locations. ST expression data suffers from high noise and drop-out events; however, smoothing techniques have the promise to improve the data interpretability prior to performing downstream analyses. Single cell RNA sequencing (scRNA-seq) data similarly suffer from these limitations, and smoothing methods developed for scRNA-seq can only utilize associations in transcriptome space (also known as one-factor smoothing methods). Since they do not account for spatial relationships, these one-factor smoothing methods cannot take full advantage ST data. In this study, we present a novel two-factor smoothing technique, Spatial and Pattern Combined Smoothing (SPCS), that employs k-nearest neighbor technique to utilize information from transcriptome and spatial relationships. By performing SPCS on multiple ST slides from pancreatic ductal adenocarcinoma (PDAC), dorsolateral prefrontal cortex (DLPFC), and simulated high-grade serous ovarian cancer (HGSOC) datasets, smoothed ST slides have better separability, partition accuracy, and biological interpretability than the ones smoothed by pre-existing one-factor methods. Source code of SPCS is provided in Github (https://github.com/Usos/SPCS).

## Introduction

Mamallian tissue is highly heterogeneous with phenotypes that depend on their spatial distribution [1, 2]. Until recently, studies of tissue heterogeneity have either sacrificed spatial relationships (e.g. scRNAseq) or produced low-dimensional measurements (e.g. IHC) [3–7]. Novel spatial transcriptomics (ST) techniques allow whole transcriptome profiles to be measured while preserving spatial relationships [8, 9]. These techniques have already been profoundly useful in understanding tumor [10–12] and non-tumor tissue [13–16] heterogeneity. However, improvements in ST library preparation [17], sequencing techniques, and bioinformatic analysis pipelines [18] are still necessary and ongoing, in contrast to more establish scRNA-seq standard practice protocols [19, 20].

The most widely utilized ST technologies are based on highly multiplexed sequence barcoding, which suffers from expression noise and drop-out events [21, 22]. Barcoding-based scRNAseq data suffers from similar limitations, while plate- and in vitro transcription-based techniques such as Smart-seq2 [23] and CEL-seq2 [24], respectively, provide more representative expression profiles per cell at the cost of fewer cells measured per experiment. As a result, a multitude of techniques have been developed to impute the missing expression values and smooth the noise that come directly from the barcode-based non-spatial scRNA-seq. SAVER [25] and MAGIC [26] use sets of correlated genes and relative cell similarity in transcriptome space to impute the drop-out events and eliminate other types of expression errors via machine learning techniques. These methods are termed “one-factor methods” given that they only incorporate expression values. The smoothened expression values give more accurate representations of the true underlying RNA abundances than the raw read counts. ST data has the advantage of providing spatial relationships that can be used in addition to transcriptomic similarity for smoothing, based on the assumption that nearby cells will have more similar expression profiles than distant cells.

Here we present a novel two-factor smoothing method, termed Spatial and Pattern Combined Smoothing (SPCS), specifically designed for ST data that utilizes both the associations of spatial locations in transcriptome space (expression pattern knowledge) and in Euclidean space (spatial knowledge). By performing SPCS on multiple ST slides from pancreatic ductal adenocarcinoma (PDAC), dorsolateral prefrontal cortex (DLPFC), and simulated high-grade serous ovarian cancer (HGSOC) datasets, smoothed ST slides have better separability, partition accuracy, and biological interpretability than the ones smoothed by pre-existing one-factor methods.

## Methods

### Datasets

The datasets that we use in this study including two real world ST dataset, PDAC [10] and DLPFC [27], and a simulating dataset generated from HGSOC single-cell dataset [28]. For the two real world datasets, PDAC include 10 ST slides sourced from traditional ST platform, while DLPFC is a Visium platform dataset with 12 slides. All the data in these datasets consist of two different matrices containing gene wise expressions and spatial coordinates. One matrix consists of the gene expression values for each spatial barcode hybridized from its corresponding spot on the ST slide. The other matrix contains the spatial locations in 2D space for each spot’s spatial barcode. Using these two matrices, we can generate a 2D representation of each gene’s expression value throughout the biopsied tissue section. Because these datasets sourced from different ST platform, i.e. traditional ST platform and new developed Visium platform, we can explore the influence of smoothing methods more comprehensively. A detailed summary statistics of the data we used is provided in Supplementary Table S1.

To better explore the ability of smoothing methods to deal with outlier spots, we designed a simulation experiment based on those used in the BayesSpace study [29]. Simulated ST data is based on HGSOC single-cell dataset and an immunofluorescence stained image of an ovarian cancer (OC) biopsy. In the original single-cell analysis of the HGSOC dataset, all the cells were divided into 15 clusters by the DBSCAN clustering method and annotated [28]. Considering the limited number of cells, we only used some of the slide. Ground-truth cluster labels were derived from single-cell level annotation of tumor and stroma compartments within the image. To make the simulated data reflect biology, we separated the slide into four clusters: intratumor (including dendritic and fibroblast cells), stroma (corresponding to macrophage cells) and two tumor clusters (associated with two different maliganent cell clusters). Detailed information about ground truth clusters is provided in Supplementary Table S2. To test the ability of different smoothing methods to find outlier spots, we randomly mixed 5% of other cell types as perturbation for each spatial cluster. We generated 10 sets of simulating data in the simulation analysis.

### Spatial and Pattern Combined Smoothing (SPCS) of Spatial Transcriptomic Expression

For each spot on an ST slide, there exists not only the gene expression but also their spatial positions. This means we can improve the quality of the expression values within each specific spot using the relative similarity to the other spots based on both expression pattern and spatial location on the ST slide. To achieve this goal, we propose a KNN-based method, Spatial and Pattern Combined Smoothing (SPCS), to perform the smoothing and padding. We display the procedure of SPCS method in Figure 1.

**Figure 1.**
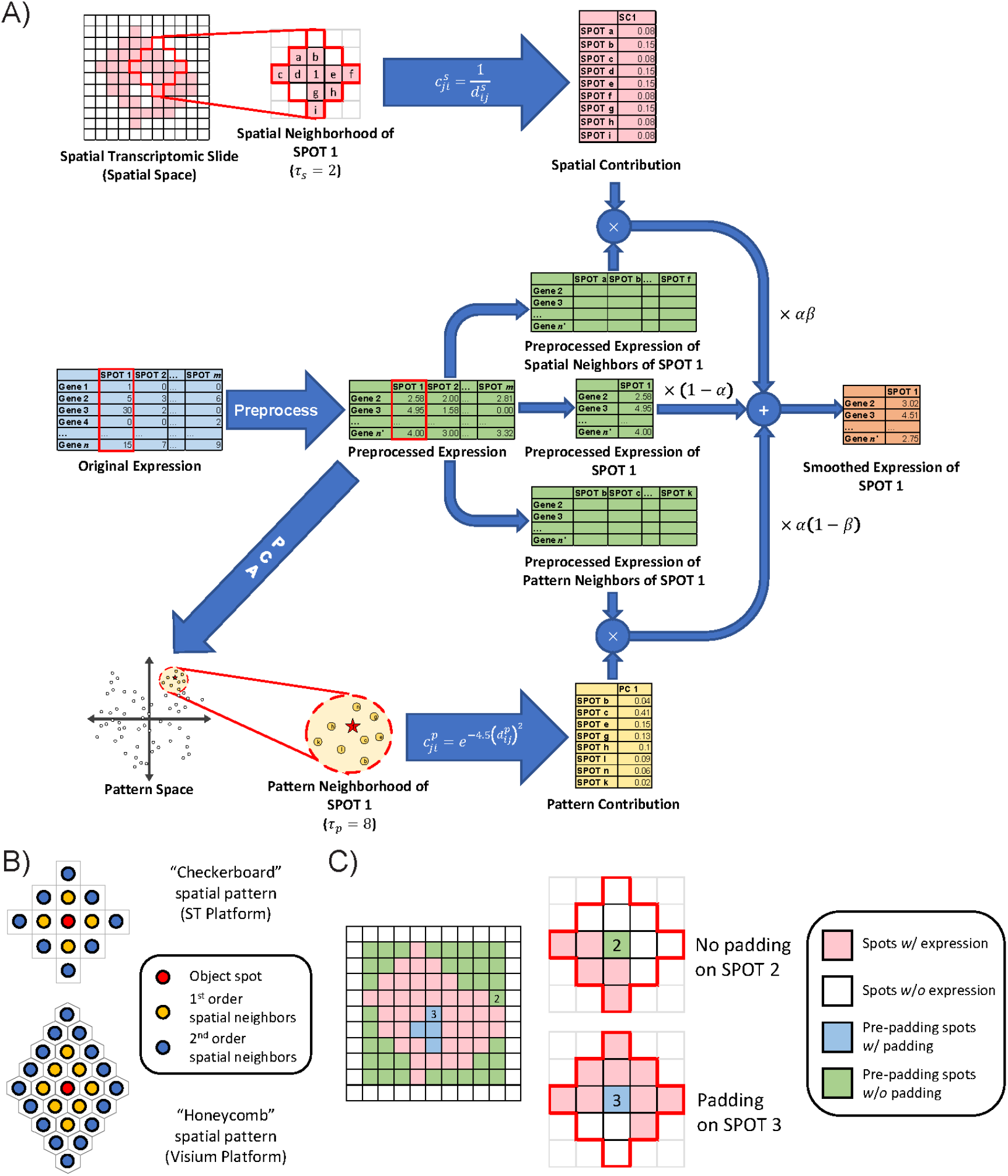
Workflow of our proposed SPCS smoothing method. (a) An example of smoothing. In this sample, SPOT 1 is the object spot that is going to be smoothed. For ST slides, preprocessing steps normalize the expressions and filter out genes with zero expression in most spots. Spatial neighborhood is second order neighborhood of SPOT 1 with nine spots. Spots where all gene expression is 0 are treated as non-tissue reigons and excluded from spatial neighborhood. Pattern neighborhood consists of the top eight spots with most similar expression pattern to SPOT1. Both spatial and pattern contributions are normalized which means total contribution of all neighbors is equal to 1. Smoothing is performed by integrating parameter- and contribution-weighted expression of the object spot itself and both it’s spatial and pattern neighbors. (b) Shape of second order spatial neighborhood for ST and Visium platforms. Spatial neighborhood is determined by Manhattan distance. Spots with same Manhattan distance to object spot belong to same order of spatial neighborhood. Due to the different geometries of spots, the shape of spatial neighborhood could be different in different platforms. (c) Padding strategy of SPCS. A missing spot will not be padded unless there are more than 50% of unmissing spots inside its spatial neighborhood.

In our method, we obtained the smoothed expression of each spot by integrating the contribution-weighted expression of its pattern and spatial neighbors. Let *X_*i*_* be a vector of gene expression values for spot *i*, smoothed expression 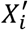 can be calculated by:

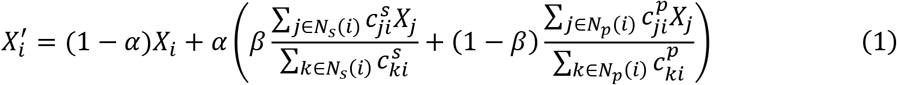

In eq. (1), there are two ratio parameters *α* and *β*. *α* is used to regularize the ratio of original expression to the corrected expression, which avoids the expression of the object spot becoming over-smoothed. *β* is used to adjust the ratio of spatial-based to pattern-based smoothing for different applications. *N_s_*(*i*) and *N_p_*(*i*) are the spatial and pattern neighborhood of the object spot *i* respectively. The size of neighborhoods can be determined by parameter *τ_s_* and *τ_p_* which also should be specified in advance. 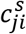 and 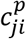 represent the spatial and pattern contributions of a corresponding neighbor spot *j* to the object spot *i*. For traditional ST platform based PDAC dataset, we set *α* = 0.6, *β* = 0.4, *τ_s_* = 2 and *τ_p_* = 16; and for the Visium platform based DLPFC and simulating datasets, we make *τ_s_* = 4 due to the larger number of spots. We will display the influence of these parameters on data seprability and discuss the selection of them in the Discussion section. In addition, SPCS will also fill the missing spots using multiple non-missing spatial neighbors. The source code of our proposed SPCS method is provided in our Github repository (https://github.com/Usos/SPCS). In the next section, we introduce the detailed mathematical definitions of neighborhood, neighborhood contribution, and our missing spot padding strategy.

### Pattern neighborhood

ST data is a type of transcriptomic data that measures gene expression patterns similar to scRNA-seq. Like some scRNA-seq data, where cells can be localized to tissue location of origin, spots which have a similar expression pattern are more likely to belong to the same region in a tissue. Therefore, smoothing the expression of a given spot using other spots with similar gene expression can improve data quality and is similar to one-factor smoothing methods [25, 26] designed for scRNA-seq. A group of the most similar spots based on the expression “pattern” of the spot can be defined as that spot’s “pattern neighborhood”. Here, we provide the explicit definition of the pattern neighborhood used by the SPCS method.

#### Definition 1

(Pattern neighborhood): *S*_*p*_ is the gene expression pattern space of an ST slide, *i*, *j*, *k* ∈ *S*_*p*_ are different spots, 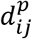 and 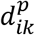 are pattern distance between spots *i*, *j* and *i*, *k* respectively. *N_p_*(*i*) is the *τ*_*p*_ pattern neighborhood of spot *i* if |*N*_*p*_(*i*)| = *τ*_*p*_, ∀*j* ∈ *N_p_*(*i*), ∀*k* ∈ *S*_*p*_ − (*N*_*p*_(*i*) *⋃* {*i*}) s.t. 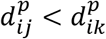.

For gene expression data, the overall shapes of gene expression patterns are of greater interest than the individual magnitudes of each feature [30]. Hence, we used Pearson correlation distance to measure pattern distance between different spots. Let *ρ*_*ij*_ represent Pearson correlation coefficient (PCC) of coordinate of spots *i* and *j* in pattern space, Pearson correlation distance *d*_*ij*_ of spots *i* and *j* is [31]:

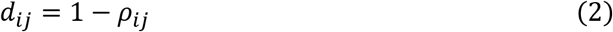

In ST data, some genes are expressed at identical or near-identical levels that lack the variance to establish an accurate pattern neighborhood. Therefore, we used principal component analysis (PCA) [32] to transform the expression of spots into a 10-dimensional principal component (PC) space before smoothing. These uncorrelated components with the largest variance from our PCA are considered the gene expression pattern space.

### Spatial neighborhood

In contrast to scRNA-seq data, ST data provide the spatial position for each spot in the slide. Regions in proximity on histopathology slides are more likely to be the same tissue type. Aside from the pattern associations between spots, we can also use spatial associations to smooth the expression as a second factor. We named the group of spots that are spatially near a given spot the “spatial neighborhood” of that spot, which is defined explicitly below.

#### Definition 2

(Spatial neighborhood): *S* represents the set of spatial location indices of an ST slide, *i*, *j* ∈ *S*, 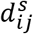 is the spatial distance between spot *i* and *j*. *N_s_*(*i*) is the *τ*_*s*_ spatial neighborhood of spot *i*, if ∀*j* ∈ *N_s_*(*i*) s.t. 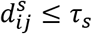.

ST spots are spatially distributed in a checkerboard or honeycomb pattern. Due to the geometric patterns inherent to ST spot layout, manhattan distance is a suitable metric to measure the spatial distance between spots inside an ST slide. Thus, we chose Manhattan distance as the spatial distance to define our spatial neighborhood. Due to the difference of spatial pattern of spots, the shape of spatial neighborhood could be different in different platform. Figure 1(b) illustrated the shape of second-order neighborhood of both traditional ST platform (checkerboard spatial pattern) and Visium platform (honeycomb spatial pattern).

### Contribution of neighbors on smoothing

Different neighbors in the spatial or pattern neighborhood, will have different impacts on the smoothing for a given spot, which we refer as “contribution”. Since the definitions of spatial and pattern distance are different, we model the corresponding contributions in different ways. The contributions of both spatial and pattern neighbors are still comparable since the range of both is [0, 1]. For spots outside the neighborhood (both spatial and pattern) of object spot, we assigned their corresponding contribution to 0, which means they have no contribution to smoothing of the object spot.

In pattern space, to better capture global gene expression patterns, we used Pearson correlation distance described in eq. (2) as the distance metric, whose range is [0, 2]. If the expression of two spots has a negative correlation, the distance based on eq. (2) will become greater than 1. However, smoothing with negative correlation spots is not performed since they are dissimilar. Therefore, we set the contribution to 0 if pattern distance between the object spot and one of its neighbors is greater than 1. We used an exponential transformation to achieve this goal. For object spot *i* and its pattern neighbor *j*, pattern contribution 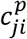 can be defined as:

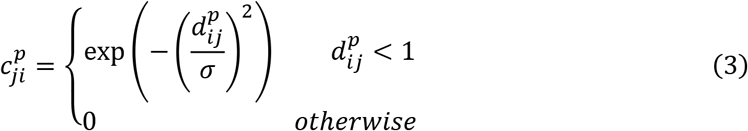

The exponential function in eq. (3) limits the range of 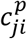 to [0, 1] and ensures 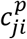 decreases as 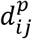 increases. *σ* in this equation is a tuning parameter that controls how the pattern contribution decays with pattern distance. When 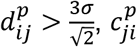 will quickly decay to 0 [33, 34]. Hence, setting *σ* to 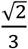 will remove the effect of negative correlation neighbors and we can simplify the eq. (3) to:

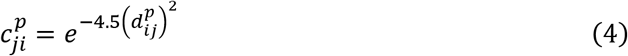

For spatial neighbors, we used Manhattan distance, an integer greater than 0, as spatial distance to measure their distance to the object spot. Since the inverse proportional function has a similar decay nature as exponential function, we used the inverse of spatial distance as the spatial contribution, i.e.:

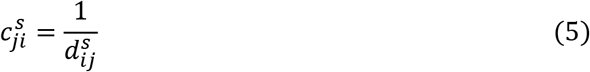

In this case, *i* is the spot being smoothed and *j* is the spatial neighbor of spot *i*.

### Padding of missing spots

For ST slides, there are two types of missing values. The first type of missing values are missing genes, which means the spot itself is located in tissue region, but some specific gene expression is missing. This is the main form of drop-out events and also frequently happens in single-cell data, which most of the smoothing methods can handle. The second type are missing spots, i.e. expression of all genes in the spot is 0. This kind of absent data is unique in ST data and cannot be padded without spatial position information, which often indicates the spot has been removed due to a quality problem. Hence, it is necessary to judge whether the missing spot need to be padded. As shown in Figure 1(c), for each blank spot in a slide, SPCS will only pad the ones whose pre-determined spatial neighborhood has more than 50% non-blank spots. This criteria ensures that boundary of the tissue will not be erroneously expanded. Since there’s no expression on missing spots at all, we estimate the expression of missing spots by their spatial neighbors only. Let *ii* become a missing spot, its expression *X_*i*_* can be estimated by:

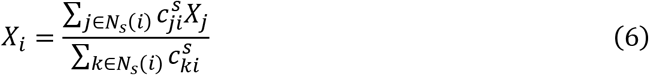

In Eq.(6), *N*_*s*_(*i*) is *τ*_*s*_ -order spatial neighborhood of spot *i*, 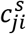 and 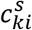 are spatial contribution of spot *j* and *k* to spot *i*. Spot padding is performed after non-missing spot smoothing. To evalulate ARI score for padding spots, we assign ground truth labels for them. The ground truth labels of the padded spots are determined by spatial contribution weighted major voting of their spatial neighbors and manually inspected by a pathology resident.

### Performance evaluation

To evaluate the effectiveness of our proposed SPCS method, we first performed SPCS and other one-factor smoothing methods (SAVER and MAGIC) on PDAC, DLPFC and simulated ST datasets. SAVER and MAGIC are two representative one-factor smoothing methods using different techniques (i.e., statistical model-based and kNN-based). Then we partitioned both smoothed and the original unsmoothed real world slides using K-medoids clustering method [35] and judged how well the clusters were separated after smoothing by internal evaluation of the unsupervised clustering. In addition, we also explored how the parameters in SPCS will influence the smoothing. Next, we performed multiple unsupervised clustering methods, including K-medoids, Louvain [36], mclust [37], BayesSpace [29] and spaGCN [38], on all the smoothed and unsmoothed slides to find out how well the clusters match the histopathological labels from the corresponding image as an external evaluation. Simulating data was also used here to detect outliers for each smoothing method. As a gene filter, we only kept genes with less than 70% zero expressed spots in our analysis. For normalization, we performed logarithmic count per million (CPM) normalization before smoothing using SPCS, SAVER, and MAGIC. The distance metric used during clustering was Pearson correlation distance as described in eq. (2). For each dataset, number of clusters is pre-determined by ground truth which is provided in Supplementary Table S1-S2. PCA was performed prior to clustering and the eigenvectors with top 20 eigenvalues were selected to reduce the dimensions of expression matrices and enhance the clustering results.

### Internal evaluation

For the internal evaluation, silhouette score was used as the evaluation indicator [39]. Silhouette score, a metric whose range is [-1,1], estimates the average distance between clusters. A greater silhouette score indicates better cluster separations. For an ST spot *i*, let (*i*) be the average distance between *i* and all the other ST spots within the same cluster and (*i*) be the smallest average distance between *i* and all the ST spots with any other clusters. The silhouette coefficient (*i*) of ST spot *i* can be expressed by:

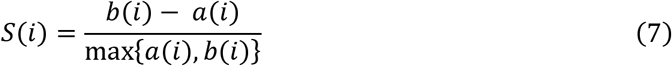

The silhouette score of an entire ST slide is the average silhouette coefficient of all spots in it.

In this work, Pearson correlation based measurment in Eq. (2) was used to calculate the dissimilarity between ST spots from the imputed ST data. Traditionally, Euclidean distance is used but here we adopted the dissimilarity metric defined in Eq. (2) since our clustering method is based on PCC distance. Padded spots are excluded in this analysis. To make the average silhouette coefficient of clustering under different smoothing methods comparable, we used the same distance matrix which was based on dimensionality reduced original unsmoothed ST expression. Hence, in our experiments, compared with unsmoothed slides, an increase or lack of change in silhouette score on smoothed slides represents the smoothing method has enhanced the original data distribution while a decrease indicates the smoothing method has corrupted the original data distribution.

### External evaluation

For the external evaluation, we obtained the histopathological labels of the ST spots. The correspondence between smoothed slides and histopathological labels was evaluated at both clustering and gene expression level. At the clustering level, the imputed ST data is clustered using multiple clustering algorithms. Technical details of the clustering methods is shown in Supplementary Table S4. Next, the concordance was evaluated between the clusters from the imputed ST data and the labels from the corresponding histopathological images. Then, the distribution of marker gene expression was compared to the locations of histopathological labels.

The Adjusted Rand Index (ARI) was used to evaluate the similarity between the clustering results from the imputed ST data and the histopathological labels [40]. For clusters from the imputed ST data {***I***_1_, ***I***_2_, ***I***_3_, ***I***_4_} and the histopathological categories of ST spots {***H***_1_, ***H***_2_, ***H***_3_, ***H***_4_}, we denoted *n*_*ij*_ as the number of ST spots that are both in ***I***_*i*_ and ***H***_*j*_. The ARI is defined as:

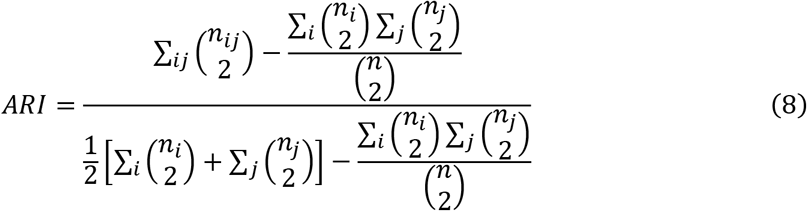

where *n*_*i*_ is the number of ST spots in ***I***_*i*_ and *n*_*j*_ is the number of ST spots in ***H***_*j*_. A higher ARI value indicates that the imputed ST clusters and the histopathological labels are more similar. In this analysis, padded spots unable to be clustered with the ground truth will be treated as an error cluster.

In the marker gene evaluation with the PDAC dataset, two marker genes, *PRSS1* and *TM4SF1*, were used to compare their expression distribution spatially. Both genes are protein-coding genes. *PRSS1* encodes a trypsinogen, which is often highly expressed in normal pancreatic tissues while *TM4SF1* is a common proto-oncogene and highly expressed in pancreatic cancer among other malignancies [41, 42]. High expression of *TM4SF1* in the cancerous regions of the PDAC dataset has been detected in previous research [10]. For the DLPFC dataset, three other marker genes, *MOBP*, *PCP4* and *SNAP25*, were used in the analysis. Previous research has reported that these genes can delineate different cortical layers [27].The expression distribution of these representative genes can stratify the histopathological regions and can be used to measure the accuracy of that partition.

### Biological analysis

To examine how our algorithm aids in informing the biology of a ST sample, we identified differentially expressed genes (DEGs) and performed gene ontology enrichment analysis (GOEA). DEGs were identified in the PDAC slides by comparing two groups of spots with *TM4SF1*-high (neoplastic tissue) and low expression (non-neoplastic tissue). To define the two groups, we first linearly transformed the expression values of *TM4SF1* between [0, 1] by dividing each value by the maximum expression value. The spots on each slide were split into two groups with one having a transformed expression value less than 0.7 (*TM4SF1* under-expressed) and the other greater (*TM4SF1* over-expressed). The cutoff 0.7 was chosen as it can reflect the boundary of the cancer region accurately.

A Kruskal-Wallis test was performed between the over- and under-expressed groups and the p-values were adjusted using the Benjamini-Hochberg method to account for the multiple comparisons [43]. Only genes with an adjusted p-value less than 0.05 were included. In addition, we used logarithmic foldchange (logFC) to determine the up-regulated and down-regulated events for genes. Genes with a logFC greater than 1 were considered up-regulated, and down-regulated if the logFC were less than -1.

GOEA was performed on the DEGs using *g:Profiler* [44]. The significance of enriched terms were tested by cumulative hypergeometric test and p-values were corrected by *g:SCS* method [45]. Only terms with an adjusted p-value below 0.05 were reported. All data sources offered by *g:Profiler* were used including Gene Ontology (GO) [46], Reactome [47], KEGG [48], WikiPathways [49], Transfac [50], CORUM [51], Human Protein Atlas [52], and the Human Phenotype Ontology [53]. Heatmaps for each sample were then generated to compare the terms found by each smoothing algorithm.

The DLPFC slides were processed slightly differently than the PDAC slides. To find the differentially expressed genes, one layer of the cortex was compared with the other six layers. For example, the expression values for all the sports in Layer 1 (L1) were compared to the expression values for the spots in layers L2, L3, L4, L5, L6, and WM (white matter). A Kruskal-Wallis test was performed between one layer of the cortex and all other layers, p-values corrected using the Benjamini-Hochberg method, and logFC calculated. This resulted in 7 sets of comparisons. The same cuttoff of p-value less than 0.05 and logFC greater than one were used. The significant DEGs were then compared to a set of DLPFC layer marker genes published by Zeng et al. [54]. Enrichment analysis was performed using *g:Profiler* in the same way as the PDAC slides were processed.

## Results

In this study, we applied our two-factor smoothing method SPCS and two state-of-the-art one-factor smoothing methods (MAGIC and SAVER) to smooth ST slides. To compare the performance of different smoothing methods, we evaluated the quality of generated clusters after performing K-Medoids unsupervised clustering on both smoothed and unsmoothed expression. The generated clusters and the distribution of marker gene expression were compared with pre-labeled histopathological partitions to check if the smoothed expression more accurately reflected the pathology features of the corresponding images. In addition, DEGs were identified in different regions with respect to the marker gene *TM4SF1* expression level and GOEA was performed to reveal the biological meaning from the smoothed data.

### Internal Evaluation

The silhouette score indicates average distance between clusters in a slide. Since we used the same unsmoothed distance matrix in the different smoothing methods, the silhouette score reflects how well the smoothing methods kept the original data distribution. For different smoothing methods, a greater silhouette score than an unsmoothed slide usually represents better separability of the smoothed expressions compared to the original data distribution. In contrast, a decreased silhouette score indicates the smoothing method has changed the original data distribution making it less seperable. After clustering the spots into corresponding clusters, we calculated the silhouette score of ten PDAC slides and eight DLPFC slides with seven clusters for each smoothing method in Figure 2(a) and Figure S1. In most of the slides from these two datasets, SPCS and MAGIC got similar or even greater silhouette score to the unsmoothed slide while SAVER got a significantly lower score. In addition, SPCS had the greatest average silhouette score over the other smoothing methods, even slightly higher than the unsmoothed slide. This result indicates that kNN-based smoothing generally can keep the charactristics of original data distribution resulting in better data separablity.

**Figure 2.**
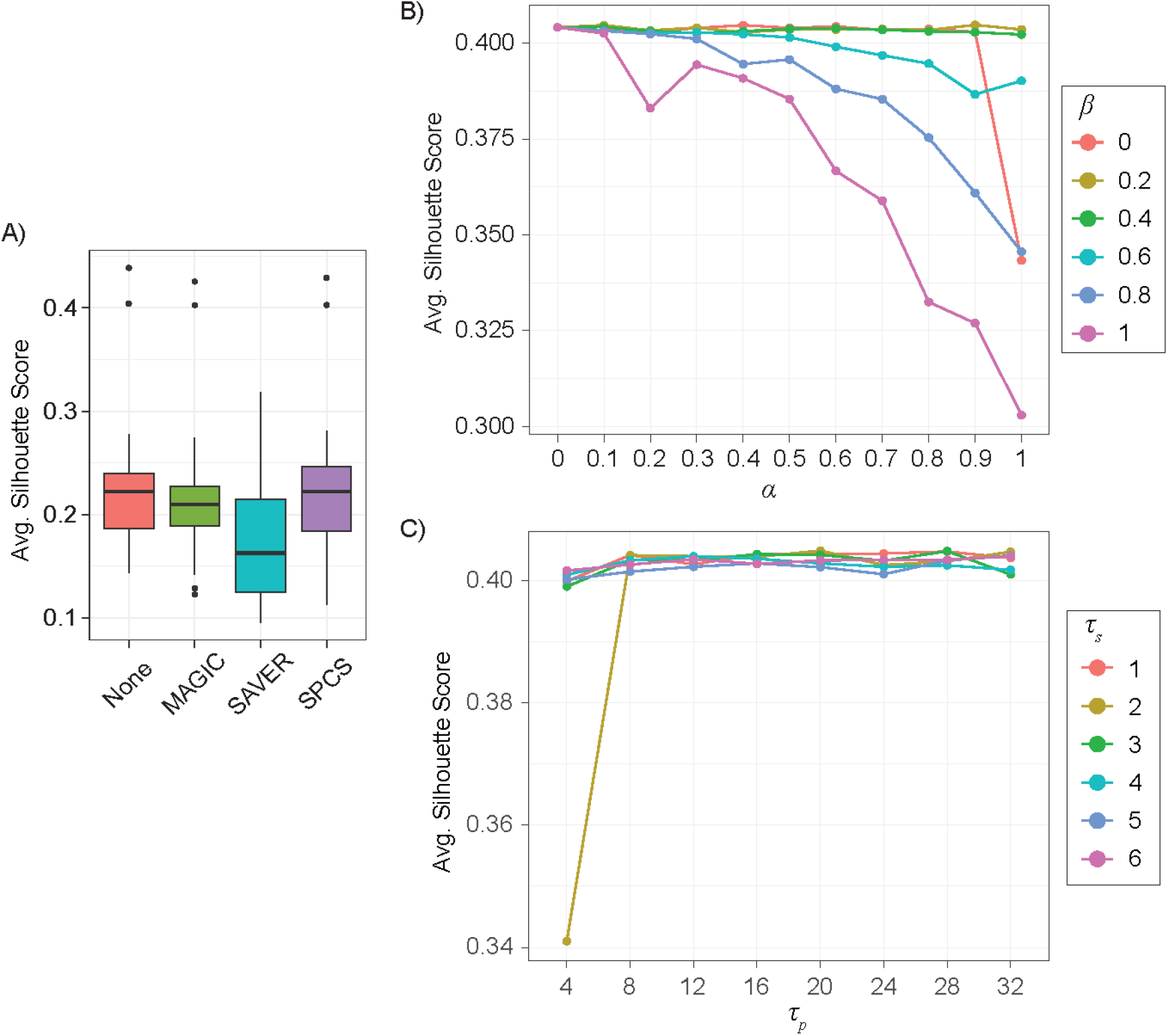
Influence of smoothing on data separability and data distribution. (a) Box plot of average silhouette score of ten PDAC samples and eight 7-layer DLPFC samples without smoothing and with smoothing by different methods (MAGIC, SAVER and SPCS). (b) Influence of parameters *α* and *β* on average silhouette score for SPCS smoothed PDACA1 slide when *τ_p_* = 16, *τ_s_* = 2. (c) Influence of parameters *τ*_*p*_ and *τ*_*s*_ on average silhouette score for SPCS smoothed PDACA1 slide when *α* = 0.6, *β* = 0.4.

Figure 2 (b)-(c) illustrated how the parameters influenced the data separability of SPCS smoothed PDACA1 slide. For the two ratio parameters *α* and *β*, while *β* ≤ 0.4, silhouette score were not affected much by the changing of *α*, but the result became more sensentive to *α* once *β* is large. This result indicated that pattern based smoothing can help keep the original data distribution. In addition, the results reveal that the data distribution of spatial neighbors is different from the pattern neighbors. For the neighborhood size parameters, the results show that *τ_s_* and *τ_p_*, have no significant influence on data separability. However, while *τ_s_* = 2 and *τ_p_* = 4, silhouette score is significantly lower than other parameter combinations, which indicates neighborhood size parameters also should be well tuned according to the dataset to avoid unexpected results.

### External Evaluation

In the external evaluation, different smoothing methods were evaluated based on the consistency of unsupervised clusters with histopathological labels. The result of PDACA1 alone was shown in Figure 3 since the histopathological labels were not available for other PDAC slides. This slide can be well clustered by K-Medoids clustering even without smoothing. When examining the results in detail, Figure 3(b) shows that most ST spots in the cancer cells and desmoplasia region were separated from other clusters. Most ST spots in the duct epithelium region were also well spearated, with slight mixing of the interstitium and the normal pancreatic tissue regions. It is worth mentioning that by including spatial position information, SPCS can pad missing spots, which other one-factor smoothing method cannot do. Figure 3(c, d) show the influence of different smoothing methods on marker genes expression. Compared with other smoothing methods, SPCS generated better marker gene contrast between distinctive histopathological areas due to two reasons. First, marker gene expressions showed fewer drop-outs and better reflected expression patterns across different tissue regions with SAVER and SPCS (MAGIC does not impute the missing expressions). Second and in contrast to SAVER, SPCS imputed the missing values and kept the spatial distribution of non-missing values stable.

**Figure 3.**
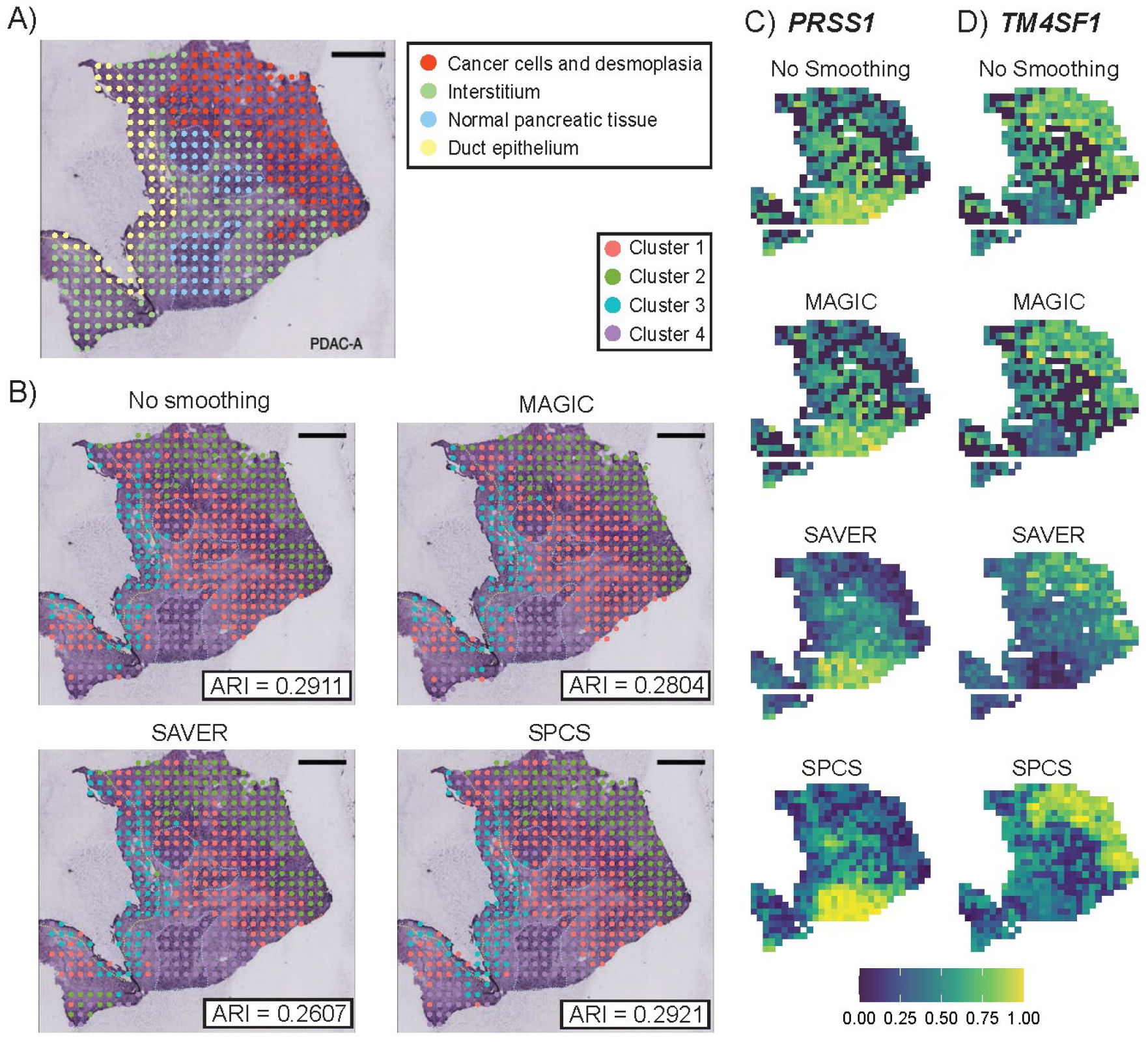
Influence of smoothing on clustering accuracy in PDACA1. (a) Original ST slide of PDACA1 and its histopathological partitions. (b) Results of K-Medoids clustering on expression smoothed by different methods. ARI score, which reflects the correlation between clusters and histopathological labels, is marked at bottom-right corner of each figure. Clusters are ordered by their size. Heatmap of smoothed expression of two marker genes, (c) *PRSS1* and (d) *TM4SF1*, are also shown. For demonstration purposes, expressions of genes are linear transformed into range of 0 to 1 as normalization.

Figure 4 illustrates external evaluation results from 12 slides in DLPFC dataset. Figure 4(a) shows the influence of smoothing methods on different clustering methods. To make the evaluation comprehensive, we chose three commonly used general clustering methods (i.e. k-Medoids, Louvain and mclust) and two state-of-the-art clustering methods developed specifically for ST slides (i.e BayesSpace and SpaGCN). The results revealed that smoothing achieved a higher ARI score using various clustering methods and SPCS outperform other one-factor smoothing methods for every clustering method. When combining SPCS with Louvain or mclust clustering methods on the DLPFC dataset, we got an ARI score near BayesSpace, indicating that the combination of SPCS with general clustering methods improves the performance of spatial clustering on ST slides. In addition, combining SPCS and BayesSpace got the highest ARI score among all the combinations. Due to the difference of preprocessing steps, the results of SpaGCN clustering were moderately lower than the original publication [38], which indicates that hyperparameter settings of SpaGCN is sensitive to preprocessing steps and we hope to optimize this in the future. Figure 4(b) and Figure S2(a) illustrated clustering ground truth and BayesSpace clustering results combined with different smoothing methods on slide 151675 and 151673. Compared with other one-factor smoothing or without smoothing, SPCS can achieve more clear and accurate boundaries between different regions, which contributes to a higher ARI score. In addition, smaller spatial neighborhood (*τ_s_*) for SPCS can help to capture long narrow regions in the slide but may cause over clustering in thicker regions with similar length and width. Similar to the PDAC dataset, SPCS also helps to obtain a better marker gene contrast between distinctive cortical layers in DLPFC dataset as shown in Figure S2(b-d).

**Figure 4.**
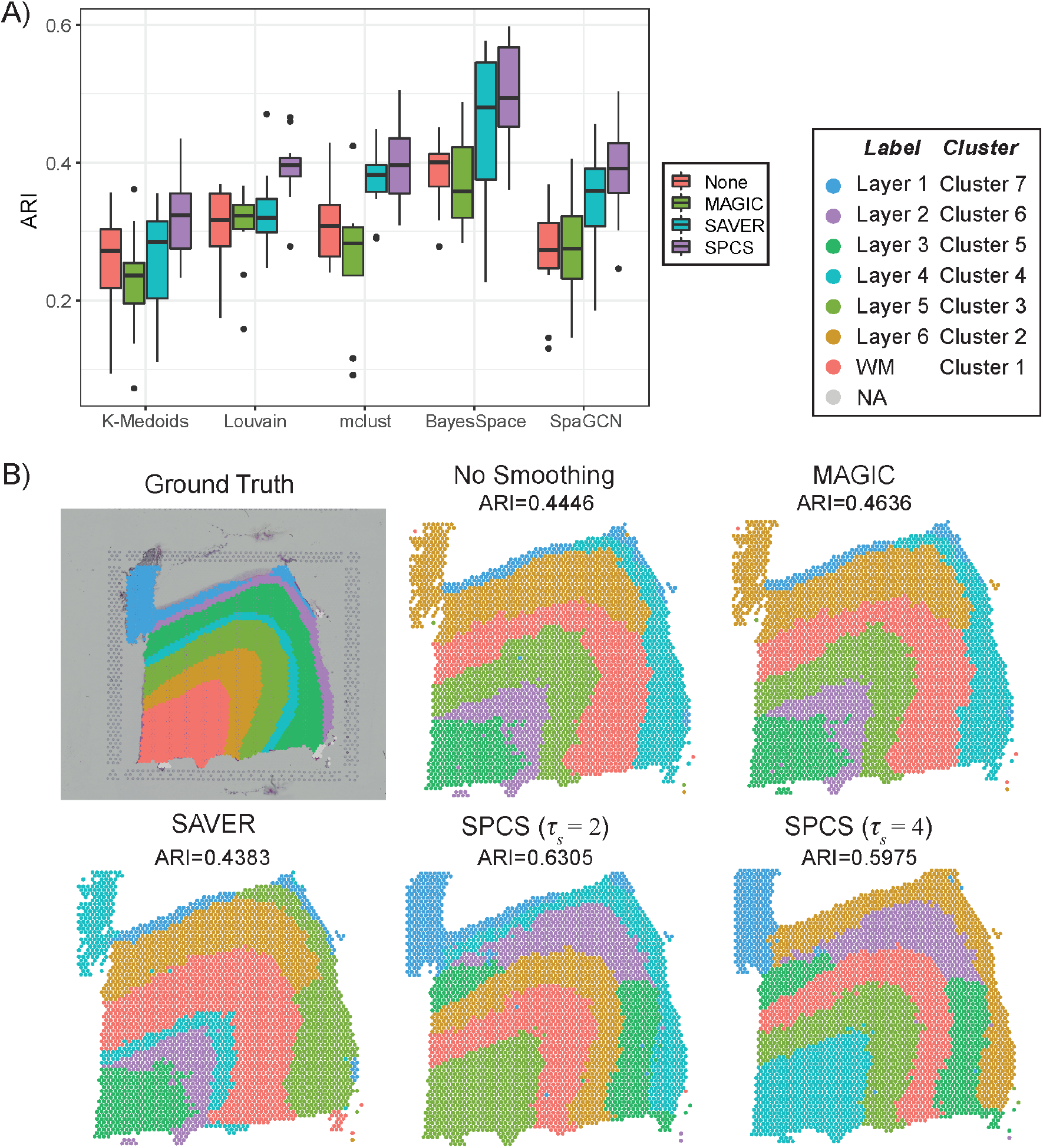
Influence of smoothing on clustering accuracy in DLPFC dataset. (a) Box plot of ARI score on various combination of smoothing (No smoothing, MAGIC, SAVER and SPCS) and clustering methods (k-medoids, Louvain, mclust, BayesSpace and SpaGCN) in 12 DLPFC samples. (b) Ground truth label and BayesSpace clustering results of smoothed and unsmoothed sample 151675. Clusters are ordered by their size.

We also performed simulation analysis to test the clustering accuracy for different smoothing methods. Louvain and BayesSpace, the two best general and ST dedicated clustering methods in the DLPFC experiment, were used in this analysis. The clustering ground truth is shown in Figure 5(a). For the 10 simulated slides, Figure 5(b) illustrated distribution of ARI score of each combination of smoothing and clustering method. In general, since the original single cell dataset was well clustered, every combination of smoothing and clustering methods got high ARI scores in this experiment. With Louvain clustering, there was only a small difference in ARI scores on different slides. For different smoothing methods, SPCS gets a slightly higher average ARI score which indicates that spatial information helps general clustering methods to cluster ST slides. However, Louvain seperates the spots into five clusters instead of four, which means Louvain tends to cluster the spots according to cell types. For the BayesSpace experiments, the ARI scores were lower than Louvain on average and greater in variance. To better review the higher variance of BayesSpace, we illustrated BayesSpace clustering results of two simulating slides in Figure 5(c) and (d). Obviously, BayesSpace tends to merge small clusters into nearby larger ones which leads to oversmoothing in the simulated dataset, and smoothing with SAVER and SPCS will aggravate this problem. Overall all methods performed well on the simulated data (ARI > 0.85) but it is worth noting that SPCS has higher potential ARI as evaluated by the 75^th^ quantile. This is surprising considering the other smoothing methods are specifically designed for scRNA-seq data from which these simulated ST slides are generated.

**Figure 5.**
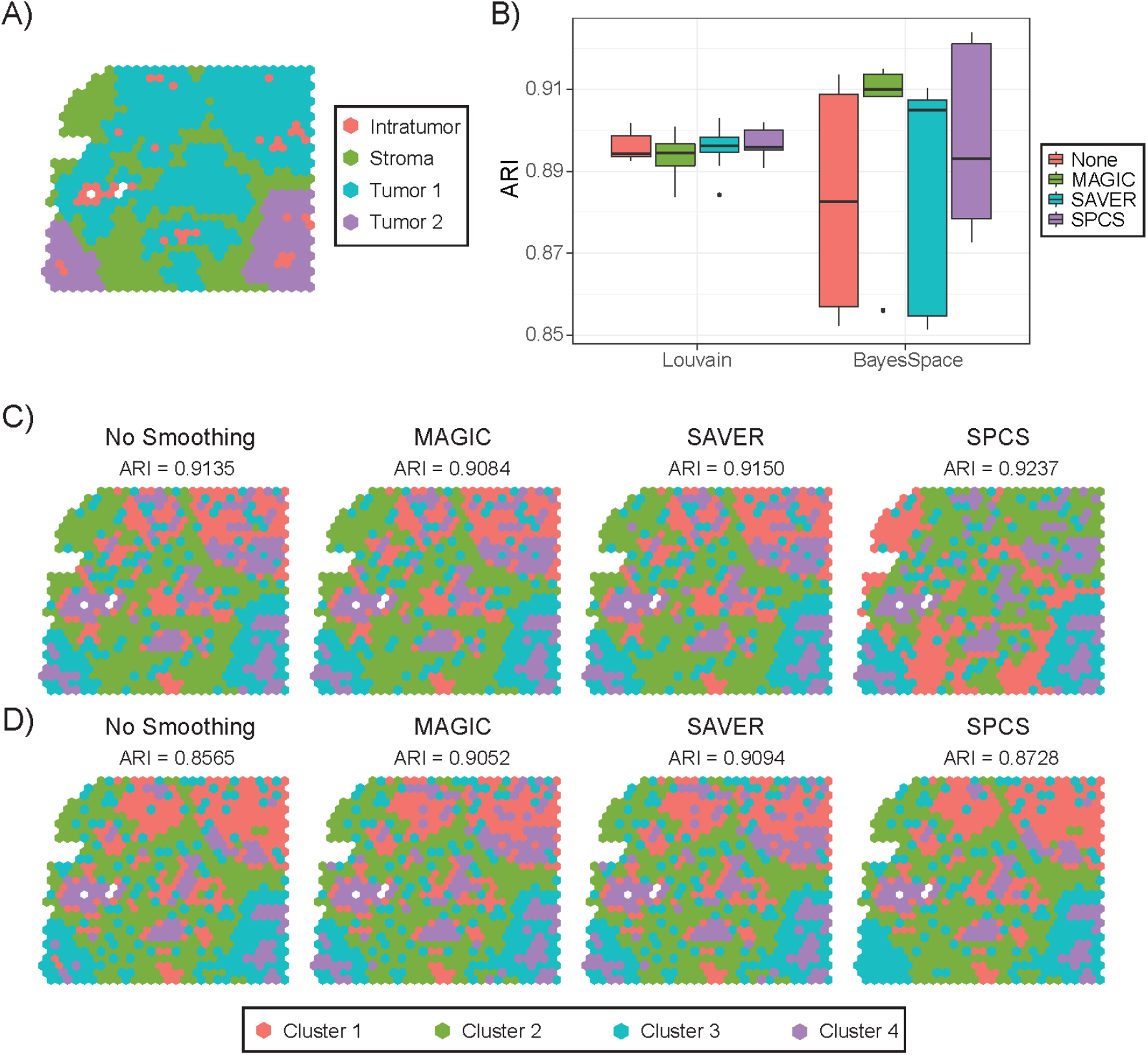
Influence of smoothing on clustering accuracy in HGSOC simulating dataset. (a) Ground truth labels of a simulated ST slide. (b) Box plot of ARI score on Louvain and BayesSpace clustered 10 smoothed simulated ST slides. (c) and (d) illustrate BayesSpace clustering results of two of the simulated ST slides. Clusters are ordered by their size.

### Biological Analysis

The biological interpretability of the smoothed results was compared between different smoothing methods. Comparing between *TM4SF1 over- and under-expressed* regions for the PDAC slides, the significant DEG numbers were displayed in Figure 6(a). There were no DEGs found in any smoothed or unsmoothed slides of PDACB2 and PDACG because they lacked *TM4SF1* while a higher number of DEGs were detected by SPCS in six out of the rest eight ST slides. Correspondingly, the number of enriched GO terms were also more from SPCS than the other methods, as shown by Figure 6(b), which are further examined below.

**Figure 6.**
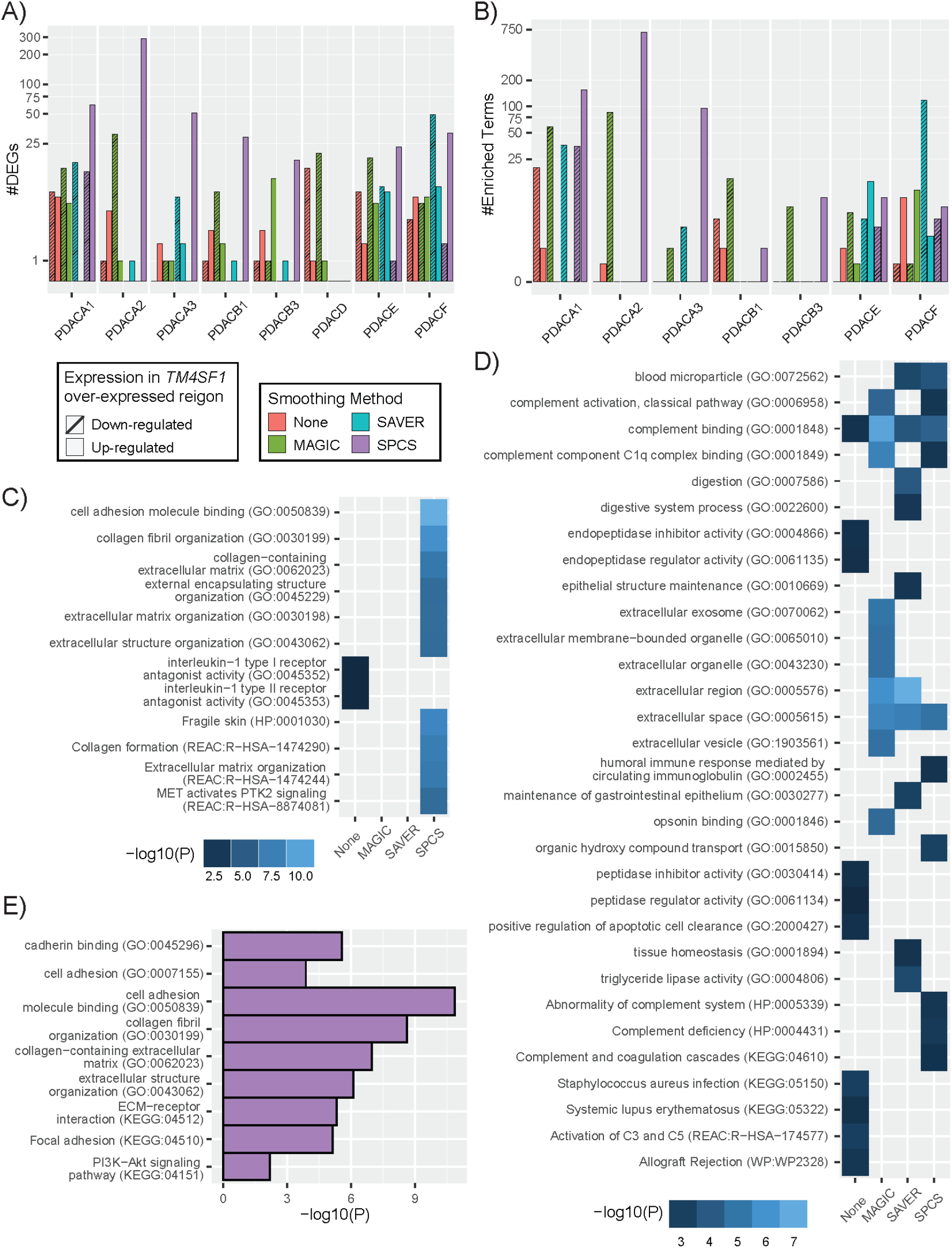
Biological analysis of unsmoothed and different methods (MAGIC, SAVER and SPCS) smoothed PDAC slides. (a) Number of DEGs identified in each slide and (b) the number of GO terms found from GOEA in each slide shown on a log10 scale. DEGs which are down-regulated in *TM4SF1* over-expressed region and their corresponding GOEA terms are marked as slash filled texture. The ten most significantly enriched terms from each smoothing method found in the (c) up-regulated DEGs and (d) down-regulated DEGs groups in *TM4SF1* over-expressed region for PDACA1 slide are shown, shaded by - log10(p-value). (e) GO terms enriched in SPCS smoothed slides from up-regulated DEGs that have previously been found in PDAC.

The top ten most significant GO terms found in slide PDACA1 for each algorithm are shown in Figure 6(c) for the up-regulated DEGs and Figure 6(d) for the down-regulated DEGs. More than ten terms are shown in the heatmap because most GO terms were not shared between smoothing methods. Without smoothing the slides, the up-regulated DEGs identified are related to interleukin-1. Instead, the terms found by SPCS smoothing are related primarily to cell adhesion, extracellular matrix (ECM) organization and MET activates PTK2 signaling. The GO terms from all smoothed and unsmoothed up-regulated slides can be seen in Figure S3. *g:Profiler* could not find any enriched terms for slides PDACB2, PDACD, and PDACG. Up-regulated enriched terms for all SPCS slides except PDACB1 appear similar to PDACA1. The results for the other methods had a small number of terms which do not seem related to PDAC, except for slide PDACF which contained cell adhesion and ECM terms for the no smoothing and MAGIC.

The enriched GO terms for the down-regulated DEGs (Figure 6(d)) tell a different story. Without smoothing, GO terms related to digestion as well as infection and autoimmune related pathways and complement cascade components (C3, C5) were identified. GO terms related to digestion were seen with SAVER as well. After performing smoothing, we found many more GO terms related to the ECM, more complement binding, as well as apoptosis regulation, but the infection and autoimmune-related pathways were absent. Applying MAGIC and SPCS also helped to find other complement cascade components such as C1q complex binding. The other slides showed are in Figure S3 and have similar results to the terms described above.

Figure 6(e) contains terms enriched in the SPCS smoothed data from up-regulated DEGs that have previously been reported in PDAC [55–58]. These terms, such as cell adhesion, cadherin binding, PI3K-Akt signaling pathway and focal adhesion, are all significantly enriched in DEGs from SPCS-smoothed data (albeit not among the top-ten enriched terms), but were absent from unsmoothed data, reflecting the enhancement of biological interpretability by performing our novel SPCS smoothing method.

The results of the biological analysis of the DLPFC slides are shown in Figure S4. The significant DEGs are shown in Figure S4(a) and show a similar number of DEGs for each algorithm. When comparing the DEGs to those reported by Zeng et al. [54], a similar number of genes were shared for each smoothing method as shown in Figure S4(b). These genes include *MOBP* (an oligodendroctye and white matter marker), *PCP4* (associated with L5 and L6) [27, 54], and *GFAP* (associated with astrocytes and many neurological disorders) [59]. In addition, comparing with other smoothing methods, SPCS helped to discover five DEGs (i.e., *CADM4*, *ELOVL5*, *AGR2*, *LGALS3BP* and *SCARB2*) which the authors have not found reported in the DLPFC. These genes were found in the white matter layers of the brain. The expression of *CADM4* and *SCARB2* in sample 151673 are shown in Figure S4(c) and Figure S4(d), respectively.

## Discussion

### Importance of smoothing on ST data

ST data is based on highly multiplexed sequence analysis where barcodes are used to split the sequenced reads into their respective tissue locations. However, this type of sequencing suffers from high noise and drop-out events. To keep enough genes to perform biological analysis, we set a relaxed filtering threshold (less than 70% zero expressed spots) to filter out non-expressed genes in preprocessing steps. Even with this rather relaxed threshold, less than 10% of genes in slides of both PDAC and DLPFC datasets were left, which indicated that drop-out events are highly frequent in ST datasets. By visualizing the expression distribution of two marker genes, *PRSS1* and *TM4SF1* in Figure 3, the gene-level drop-out events can be easily seen, as indicated by the black-colored spots on the slides. For both genes, drop-out events frequently occurred in interstitium regions of the slide, leading to a failure cluster this area on unsmoothed ST data. In addition, there’s also entire spots missing in multiple ST slides, which can negatively influence spatial clustering. By performing smoothing methods, the missing and noisy expression values were controlled to a certain extent so that the partition of regions and downstream analysis are greatly improved. Therefore, smoothing is an important and necessary step for analyzing ST data.

### SPCS Improves Data Quality

Compared with unsmoothed data, smoothing improves data quality. We have demonstrated that various smoothing algorithms increase the separability and partition accuracy of ST spots. Moreover, by performing internal and external evaluations, we confirm that SPCS-smoothed data show better quality as compared with the two existing one-factor smoothing methods MAGIC and SAVER. For internal evaluation, from the results shown in Figure 2, SPCS smoothing method produces greater silhouette scores than MAGIC and SAVER, which means the ST data smoothed by SPCS has better separability. In addition, a more similar silhouette score of SPCS smoothed and original unsmoothed data indicates that SPCS can better preserve the original data distribution which helps to keep accurate biological analysis results while improving spatial clustering accuracy.

Our external evaluation verifies the partition accuracy of smoothed data. One main objective of slide partition is to identify different histopathological regions in the slide. Hence, we measured the degree of overlap between unsupervised clusters on smoothed data and histopathological partitions using ARI. Since the spots in the same histopathological region are usually connected, incorporating spatial knowledge is expected to boost partition accuracy. Moreover, spatial knowledge can also help to detect and pad missing spots. Indeed, as expected, results in Figure 3-5 reveal SPCS method generates a higher ARI score than existing one-factor methods, which means a more accurate histopathological partition can be acquired by performing two-factor SPCS method. In addition, it is also clear that SPCS can be used before various clustering methods to improve clustering accuracy. From the marker gene analysis, SPCS method recovered the drop-out events and enhanced the expressions of marker genes in the corresponding regions. This evidence proves that SPCS can improve the accuracy of the ST spot partitions.

### SPCS Enhances Biological Interpretability

Our SPCS method identified many more differentially expressed genes than the other smoothing methods tested for most of the ST slides. *IL1RN*, *KRT7*, *LAMB3*, *LAMC2*, *NOTCH3, and S100A16* were identified as upregulated DEGs in the PDACA1 slide after SPCS processing, and each of these genes has been associated with poor survival in PDAC [56, 63]. *IL1RN* and *LAMB3* were also found using unsmoothed data and *LAMB3* was detected with MAGIC smoothed data, but the other genes were uniquely detected by SPCS. A higher number of DEGs associated with GO terms after SPCS processing than unsmoothed, SAVER, and MAGIC processing. Specifically, only slides PDACE and PDACF contained up-regulated DEGs and GO for SAVER and MAGIC; although PDACD was the only slide which had DEGs and no GO terms for the down-regulated genes. The unsmoothed up-regulated slides PDACA2, PDACB3, PDACD, and PDACE, and down-regulated slides PDACA3, PDACB3, and PDACD also had DEGs but no GO terms. In contrast, every SPCS smoothed slides that had DEGs also had GO terms. The SPCS data demonstrated a *TM4SF1* expression landscape that matched the original histopathological assignment more accurately than the other smoothing methods.

The GO terms reported in PDACA1 using SPCS related more to pancreatic cancer and the role of *TM4SF1* than the non-smoothed data and the two one-factor smoothed data. *TM4SF1* has been found to be overexpressed in PDAC and has roles in apoptosis, proliferation, and cell migration [41]. Previous work found that collagen 1 (GO:0030199) binds with DDR1 which then interacts with TM4SF1 to activate the focal adhesion kinase (FAK) [41, 60, 61]. This results in disruption of E-cadherin (GO:0045296) leading to the Wnt signaling pathway and loss of cell-cell adhesion (GO:0050839) [41, 60]. The FAK pathway also increases expression of N-cadherin (GO:0045296) which results in migration of cancer cells [41, 60]. TM4SF1 can also activate the *AKT* pathway (KEGG:04151) leading to anti-apoptotic effects and angiogenesis [41, 62]. All of these previously identified GO terms are important factors for PDAC survival and metastasis. They were also identified using SPCS-smoothed data only, which would have been missed using other smoothing methods or with the unsmoothed ST data.

The terms found through GOEA were more interpretable in the scenario of PDAC and its pathogenesis when applying SPCS smoothing. Many of the SPCS top ten terms in Figure 4(c) and previously reported terms in Figure 4(e) are similar to those found in the literature [55–58] and are consistent between slides. Without applying SPCS, some of the top terms found in Figure 4(d) are involved in typical pancreatic activity such as peptidase regulator activity (GO:0061134), digestion (GO:0007586), and triglyceride lipase activity (GO:0004806) which indicates that important PDAC-pathology related GO terms may be missed when data is not smoothed or not properly smoothed.

Identifying the DEGs in different histological regions and checking their associated GO terms can help evaluate the biological interpretability of a smoothed slide. It is worth noting that the gene ontology (i.e. gene set database) and the enrichment tool can yield different results. The simpliest example would be the use of a hypergeometric test to determine enriched gene sets opposed to gene set enrichment analysis which accounts for the significance of DEGs. The hypergeometric test has a longer history of use and is easily interpretable whereas gene set enrichment analysis is a newer approach. Furthermore, the gene sets themselves differ between databases such as KEGG and GO. There could potentially be more enriched KEGG pathways for one sample and more enriched GO terms for another. For these reasons, we used g:Profiler since it incorporates many gene set databases in the analysis and relies on the well-established hypergeometric testing approach.

Using the cortical layers in the DLPFC slides, we found a similar number of DEGs and shared genes between the smoothing methods. Most of these genes were previously reported. Due to the enhanced contrast, our proposed SPCS method helped to identify five DEGs in the white matter while the other methods did not. The Cell Adhesion Molecule 4 (*CADM4*) and scavenger receptor class B Member 2 (*SCARB2*), to the author’s knowledge, have not been identified as white matter markers in the DLPFC. Cell adhesion molecules (CAMs) like *CADM4* play an important role in myelination by oligodendrocytes. Higher expression of *CADM4* leads to many short myelin internodes that disrupt the normal myelination process [64]. *SCARB2* is a lysosomal membrane receptor for the glucocerebrosidase enzyme. It has been associated with Parkinson’s disease and Lewy Body Disease. Glucocerebrosidase degenerates sphingolipid, which is important for brain development. There is some evidence that decreases sphingolipids can lead to demyelination [65, 66]. While the authors do not intend to present *CADM4* and *SCARB2* as marker genes for white matter, we believe that SPCS can be used to help aid with this task. The enrichment analysis results for DLPFC are not shown but are similar between all smoothing methods. This is likely because the number of GO terms found is correlated with the number of DEGs. It’s possible that using a gene markers instead of layers to identify DEGs could produce different results. Given the generally clearly defined boundries of the brain, using layer data to get differential gene expression seems more appropriate.

### Determination of SPCS parameters

There are four parameters in SPCS: *τ_s_*, *τ_p_*, α and *β*. The parameters *τ_s_* and *τ_p_* are designed to adjust the size of spatial neighborhood and pattern neighborhood respectively. Including more information while performing smoothing is beneficial for a more robust result, and increasing the size of neighborhood is a good way to achieve that goal. Blindly expanding neighborhoods will incorporate some spots that are not similar to the spot being smoothed; therefore, SPCS uses contribution weighting to reduce this effect, which gives the size of both spatial and pattern neighborhoods limited influence on data separability as shown in Figure 2(e). In addition, as shown in Figure 4 (b) and (c), spatial neighborhood will also influence clustering sensitivity. A smaller spatial neighborhood (*τ_s_*) for SPCS can help to capture long narrow regions in slides but may cause over clustering in thicker regions with similar length and width. Therefore, we recommend a modest selection of these two parameters to balance the tradeoff. For most cases, *τ_p_* ≤ 16 and *τ_s_* ≤ 4 is recommended.

The parameters, α and β, are designed to balance the original expression and corrections from spatial and pattern neighbors, which has a significant effect on smoothing quality. Due to the pattern similarity between the object spot and its pattern neighbors, corrections from pattern neighbors tend to enhance the original data distribution features shown by an increased or stabilized on silhouette score. In contrast, corrections from spatial neighbors make the object spot expression consistent with its spatial neighbors. This is beneficial for spatial clustering but may change the original data distribution leading to a worse silhouette score. To keep the accuracy of spatial clustering and biological analysis simultaneously, it is important to balance the intensity of correction with the underlying expression signatures. Results in Figure 2(d) indicate that the influence of smoothing strength (*α*) on data separability is heavily reliant on the proportion of spatial and pattern correction (*β*). Hence, we recommend setting α between 0.2 to 0.8 and β less than 0.6 in most cases, and values should be selected carefully according to data distribution. In addition, as indicating in Figure 5, when combined with spatial clustering methods like BayesSpace, smaller *α* and *β* are recommended to avoid erroroneous merging of small clusters.

## Conclusion

In response to expression noise and drop-out events in barcoding-based sequencing technologies, smoothing has become an essential data processing step before performing the downstream analysis on ST data. In this paper, we proposed a novel two-factor ST data smoothing method, SPCS, which can take full advantage of both the expression patterns and the spatial patterns contained in ST data. Compared with traditional one-factor smoothing methods, SPCS improved separability, partition accuracy, and biological interpretability of ST experiments. SPCS can effectively improve ST data quality for accurate and meaningful downstream analyses. SPCS is broadly applicable on any barcoding-based ST technology.

## Key Points

- Due to the common issue of noise and drop-out events in ST data, smoothing has become a necessary step before downstream analysis on ST data.
- SPCS is a novel kNN-based two-factor smoothing method that can fully utilize both expression pattern and spatial knowledge in ST data.
- Compared with traditional expression pattern knowledge-based one-factor smoothing methods, SPCS can provide better separability, partition accuracy and biological interpretability.

## Data Availability

All of the data used for the analyses in the manuscript are freely available from their original publications.

## Funds

This work is partially supported by ACS-IRG Grant Mechanism [Grant No. 19-144-34, to T.S.J.], the National Natural Science Foundation of China [Grant No. 41876100, to X.Y.], the State Key Program of National Natural Science Foundation of China [Grant No. 61633004, to X.Y.], the Indiana University Precision Health Initiative [to K.H. and J.Z.].

## Acknowledgement

We special thanks Dr. Edward Zhao and Dr. Raphael Gottardo in Fred Hutchinson Cancer Research Center for providing information and help on simulation experiment design.

## Description of Authors

Yusong Liu is a PhD student in the College of Intelligent Systems Science and Engineering, Harbin Engineering University. His research is focus on machine learning and network analysis in bioinformatics.

Dr. Tongxin Wang is a recently graduated PhD student from the Department of Computer Science, Indiana University, who is currently employed as a Research Scientist at Facebook. His research focus is on deep learning, transfer learning, and adversarial networks.

Ben Duggan is a medical student at Indiana University School of Medicine. His research interests include bioinformatics, machine learning, and using computing to improve patient care.

Dr. Michael Sharpnack is a pathology resident in the Department of Pathology at the Univeristy of California San Francisco. His research is primarily concerned with novel immunotherapies, antigen presentation, and bioinformatics methods development as it applies to lung cancer.

Dr. Kun Huang is a Professor and Chair of the Department of Biostatistics and Health Data Science, Indiana University School of Medicine, Indiana University School of Medicine Precision Health Initiative Chair for Genomic Data Science, Director of Data Science and Informatics for the Precision Health Initiative, Associate Director of Data Science for the Indiana University Simon Comprehensive Cancer Center, and an Investigator at the Regenstrief Institute. His main research interests are in medical image analysis, multi-omics, and machine learning.

Dr. Jie Zhang is an Assistant Professor of Medical and Molecular Genetics who is also a member of the Center for Computational Biology and Bioinformatics. Her research interests are applied translational bioinformatics and systems biology for cancer and neurological disease.

Dr. Xiufen Ye is a professor of College of Intelligent Systems Science and Engineering, Harbin Engineering University, IEEE Senior Member. Her main research interest includes image processing, pattern recognition and artificial intelligent.

Dr. Travis S. Johnson is an Assistant Research Professor in the Department of Biostatistics and Health Data Science, Indiana University School of Medicine. His research interests include applied machine learning for use with high dimensional omic data as it applies to cancer and dementia research.

