## Supplemental tables and figures for "SPCS: A Spatial and Pattern Combined Smoothing Method for Spatial Transcriptomic Expression"

**Supplementary Table 1.** Summary statistics of real-world datasets used in the experiments. Number of SPCS padding spots was estimated based on corresponding setting of spatial neighborhood size ($\tau_s$) for each dataset. Filtered genes are the ones with no more than 70% zero expression spots in each sample. Number of clusters was obtained from previous research of corresponding datasets.

| Platform | Sample | #Spots | #Padded spots by SPCS | #Genes | #Filtered genes | #Clusters |
|---|---|---|---|---|---|---|
| **PDAC dataset [1]** | | | | | | |
| ST | PDACA1 | 428 | 23 | 19738 | 485 | 4 |
| | PDACA2 | 316 | 41 | | 432 | 4 |
| | PDACA3 | 288 | 1 | | 145 | 4 |
| | PDACB1 | 224 | 9 | | 676 | 4 |
| | PDACB2 | 321 | 18 | | 139 | 4 |
| | PDACB3 | 242 | 9 | | 250 | 4 |
| | PDACD | 460 | 34 | | 1552 | 4 |
| | PDACE | 359 | 4 | | 1081 | 4 |
| | PDACF | 359 | 12 | | 1642 | 4 |
| | PDACG | 288 | 4 | | 625 | 4 |
| **DLPFC dataset [2]** | | | | | | |
| Visium | 151507 | 4226 | 41 | 33538 | 1028 | 7 |
| | 151508 | 4384 | 44 | | 805 | 7 |
| | 151509 | 4789 | 35 | | 1036 | 7 |
| | 151510 | 4634 | 65 | | 973 | 7 |
| | 151669 | 3661 | 124 | | 1512 | 5 |
| | 151670 | 3498 | 123 | | 1362 | 5 |
| | 151671 | 4110 | 86 | | 1592 | 5 |
| | 151672 | 4015 | 120 | | 1463 | 5 |
| | 151673 | 3639 | 79 | | 2079 | 7 |
| | 151674 | 3673 | 96 | | 2818 | 7 |
| | 151675 | 3592 | 103 | | 1495 | 7 |
| | 151676 | 3460 | 109 | | 1662 | 7 |

**Supplementary Table 2.** Spatial ground truth setting for HGSOC simulated dataset. Single cell (SC) clusters were generated and annotated for cell type by [3]. Single cells are sampled into corresponding spatial clusters with 5% replacement by other types of cells as noise. Number of spots for each spatial cluster listed here include noise spots and the noise spots spatial cluster is the summarize of noise spots in all spatial clusters.

| Spatial cluster | Spatial cluster name | #Spots (Incl. noise) | Corresponding SC clusters | SC cell type | #Cells |
|---|---|---|---|---|---|
| 1 | Intratumor | 475 | 15 | Dendritic cell | 169 |

| | | | | | |
|---|---|---|---|---|---|
| | | | 8 | Fibroblast | 360 |
| 2 | Stroma | 1893 | 11 | Macrophage | 2535 |
| 3 | Tumor 1 | 1876 | 1 | Malignant | 3191 |
| 4 | Tumor 2 | 532 | 2 | Malignant | 1613 |
| All clusters | Noise spots | 235 | All other clusters | Various types | 1741 |

**Supplementary Table 3.** Average computing cost of SPCS method on sample PDACA1 in 10 runs. Experiment based on R 4.1.2 and Windows 10 running on Intel (R) Core (TM) i5-9400F CPU and 16GB RAM. I/O operation time is excluded.

| Steps | Time (s) | RAM (MB) |
|---|---|---|
| Detecting pattern neighbors | 4.38 | 17.27 |
| Detecting spatial neighbors | 4.52 | 17.20 |
| Calculating smoothed expression | 4.70 | 219.07 |
| Padding missing spots | 4.19 | 17.36 |
| Miscellaneous | 0.12 | 55.84 |
| **Total** | 17.91 | 326.74 |

**Supplementary Table 4.** Clustering methods used in external evaluation. For all clustering methods, PCA was performed before clustering as dimensional reduction and top 20 PCs were selected for each sample. Parameters not mentioned were set as default.

| Methods | Required input | Settings | Platform | Package |
|---|---|---|---|---|
| k-Medoids [4] | Expression matrix | As default settings. | R | ClusterR [5] |
| Louvain [6] | Expression matrix | Shared nearest-neighborhood (SNN) network was constructed by top 8 nearest neighbors; Resolution parameter was tuned manually to obtain specific number of clusters for each sample. | R | Seurat [7] |
| mclust [8] | Expression matrix | As default settings. | R | mclust |
| BayesSpace [9] | Expression matrix and spatial coordinate matrix | mclust as Initial clustering method; t-distribution as error model; 10000 MCMC iterations. | R | BayesSpace |
| SpaGCN [10] | Expression matrix, spatial coordinate matrix and histopathological image | As default settings. | Python | SpaGCN |

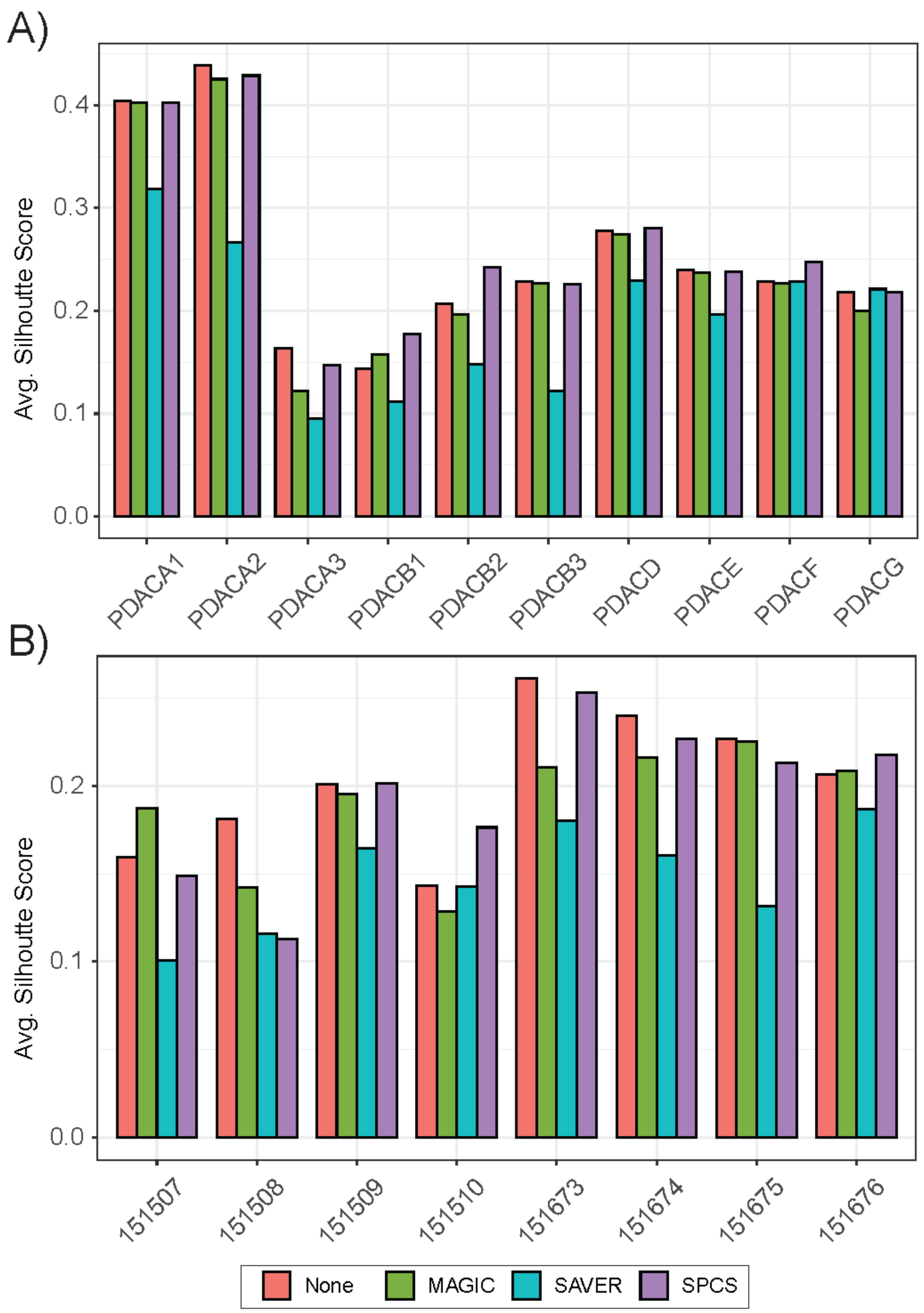


**Supplementary Figure 1.** Average Silhouette scores for (a) PDAC dataset and (b)7-layer slides in DLPFC dataset.

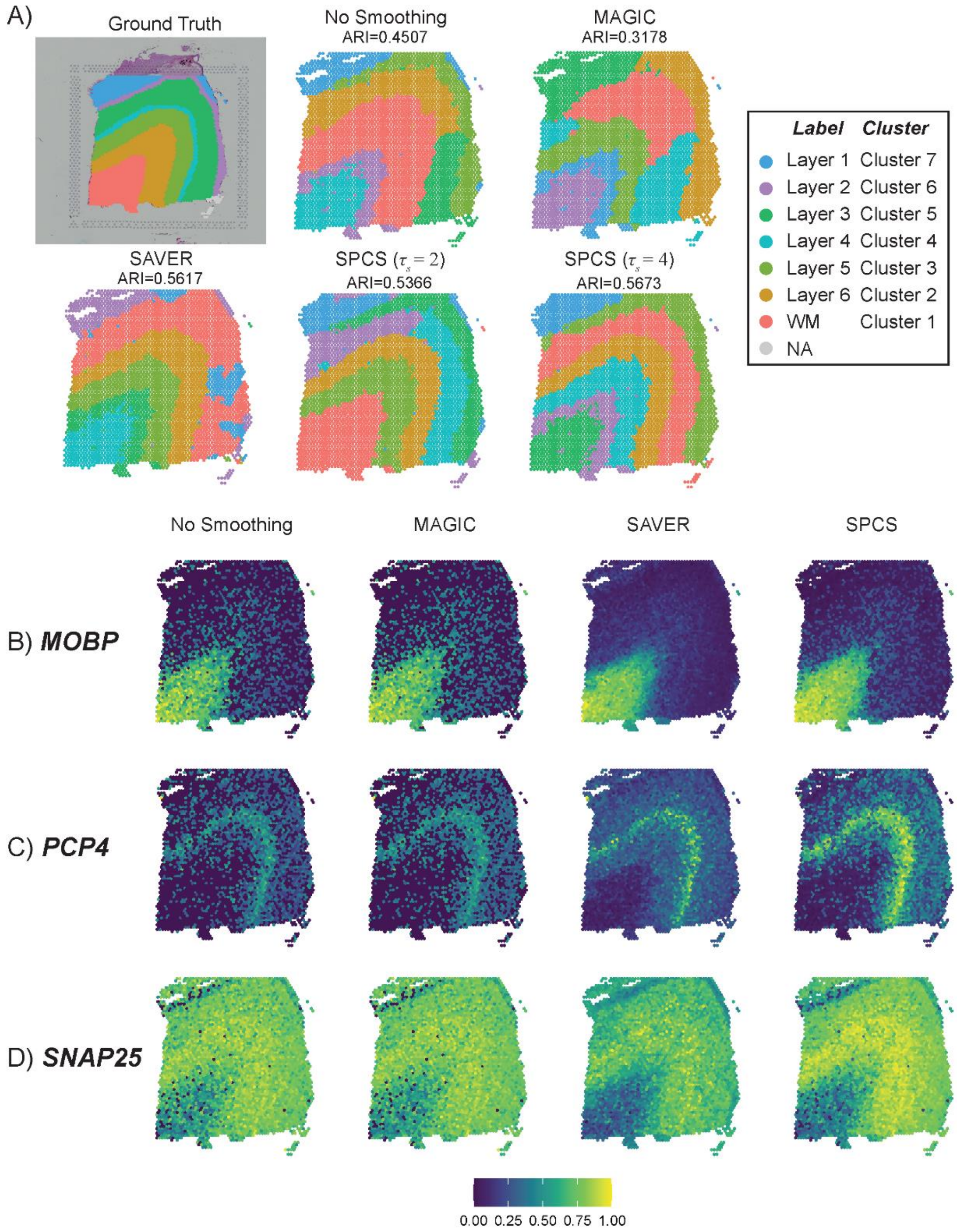


**Supplementary Figure 2.** Influence of smoothing on clustering accuracy in DLPFC sample 151673. (a) Ground truth labels and BayesSpace clustering results of smoothed and unsmoothed slide. (b)-(d) Heatmaps of unsmoothed and smoothed expressions of three marker genes, (b) *MOBP*, (c) *PCP4* and (d) *SNAP25*, which can delineate different cortical layers. For demonstration purposes, expression of genes was linearly transformed into range of 0 to 1 as normalization.

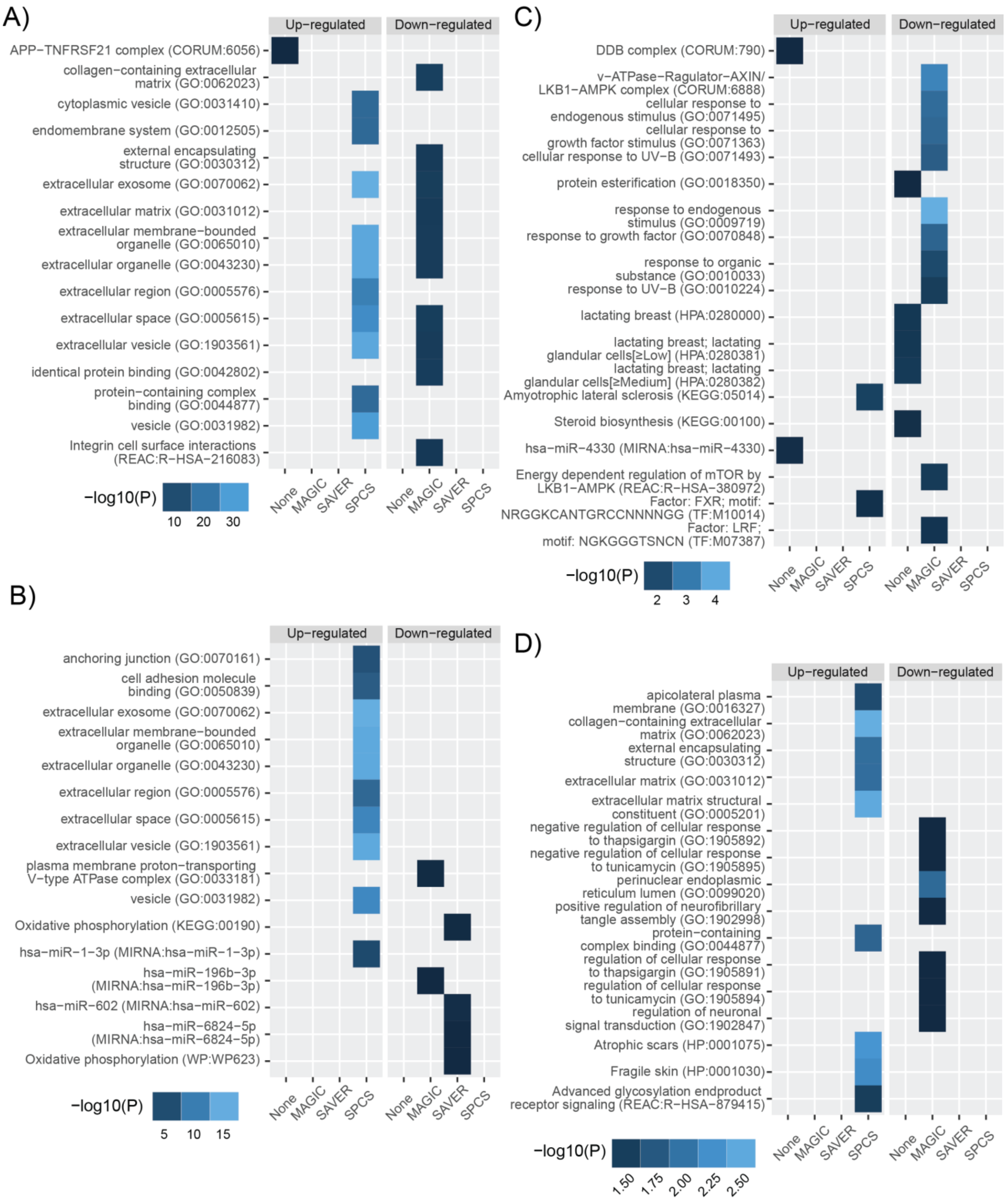
A)
Up−regulated
Down−regulated
APP−TNFRSF21 complex (CORUM:6056)
collagen−containing extracellular matrix (GO:0062023)
cytoplasmic vesicle (GO:0031410)
endomembrane system (GO:0012505)
external encapsulating structure (GO:0030312)
extracellular exosome (GO:0070062)
extracellular matrix (GO:0031012)
extracellular membrane−bounded organelle (GO:0065010)
extracellular organelle (GO:0043230)
extracellular region (GO:0005576)
extracellular space (GO:0005615)
extracellular vesicle (GO:1903561)
identical protein binding (GO:0042802)
protein−containing complex binding (GO:0044877)
vesicle (GO:0031982)
Integrin cell surface interactions (REAC:R−HSA−216083)
None
MAGIC
SAVER
SPCS
−log10(P)
10
20
30
B)
Up−regulated
Down−regulated
anchoring junction (GO:0070161)
cell adhesion molecule binding (GO:0050839)
extracellular exosome (GO:0070062)
extracellular membrane−bounded organelle (GO:0065010)
extracellular organelle (GO:0043230)
extracellular region (GO:0005576)
extracellular space (GO:0005615)
extracellular vesicle (GO:1903561)
plasma membrane proton−transporting V−type ATPase complex (GO:0033181)
vesicle (GO:0031982)
Oxidative phosphorylation (KEGG:00190)
hsa−miR−1−3p (MIRNA:hsa−miR−1−3p)
hsa−miR−196b−3p (MIRNA:hsa−miR−196b−3p)
hsa−miR−602 (MIRNA:hsa−miR−602)
hsa−miR−6824−5p (MIRNA:hsa−miR−6824−5p)
Oxidative phosphorylation (WP:WP623)
None
MAGIC
SAVER
SPCS
−log10(P)
5
10
15
C)
Up−regulated
Down−regulated
DDB complex (CORUM:790)
v−ATPase−Ragulator−AXIN/ LKB1−AMPK complex (CORUM:6888)
cellular response to endogenous stimulus (GO:0071495)
cellular response to growth factor stimulus (GO:0071363)
cellular response to UV−B (GO:0071493)
protein esterification (GO:0018350)
response to endogenous stimulus (GO:0009719)
response to growth factor (GO:0070848)
response to organic substance (GO:0010033)
response to UV−B (GO:0010224)
lactating breast (HPA:0280000)
lactating breast; lactating glandular cells[≥Low] (HPA:0280381)
lactating breast; lactating glandular cells[≥Medium] (HPA:0280382)
Amyotrophic lateral sclerosis (KEGG:05014)
Steroid biosynthesis (KEGG:00100)
hsa−miR−4330 (MIRNA:hsa−miR−4330)
Energy dependent regulation of mTOR by LKB1−AMPK (REAC:R−HSA−380972)
Factor: FXR; motif: NRGGKCANTGRCCNNNNGG (TF:M10014)
Factor: LRF; motif: NGKGGGTSNCN (TF:M07387)
None
MAGIC
SAVER
SPCS
−log10(P)
2
3
4
D)
Up−regulated
Down−regulated
apicolateral plasma membrane (GO:0016327)
collagen−containing extracellular matrix (GO:0062023)
external encapsulating structure (GO:0030312)
extracellular matrix (GO:0031012)
extracellular matrix structural constituent (GO:0005201)
negative regulation of cellular response to thapsigargin (GO:1905892)
negative regulation of cellular response to tunicamycin (GO:1905895)
perinuclear endoplasmic reticulum lumen (GO:0099020)
positive regulation of neurofibrillary tangle assembly (GO:1902998)
protein−containing complex binding (GO:0044877)
regulation of cellular response to thapsigargin (GO:1905891)
regulation of cellular response to tunicamycin (GO:1905894)
regulation of neuronal signal transduction (GO:1902847)
Atrophic scars (HP:0001075)
Fragile skin (HP:0001030)
Advanced glycosylation endproduct receptor signaling (REAC:R−HSA−879415)
None
MAGIC
SAVER
SPCS
−log10(P)
1.50
1.75
2.00
2.25
2.50

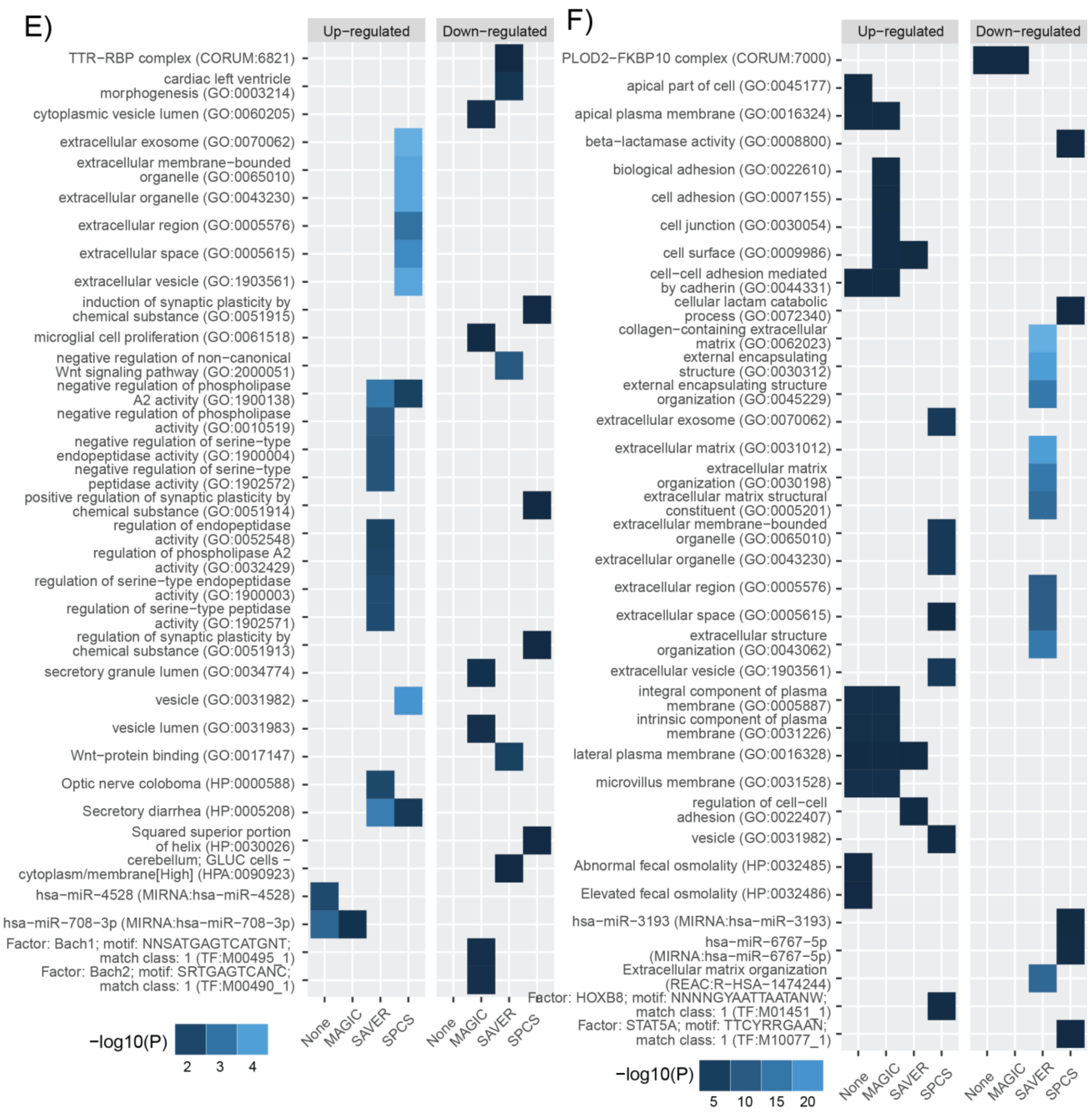


**Supplementary Figure 3.** Enriched GO terms for all smoothed and unsmoothed ST slides: (a) PDACA2 (b) PDACA3 (c) PDACB1 (d) PDACB3 (e) PDACE and (f) PDACF. The top-ten most significant GO terms are reported from each smoothing method. Results here are more than ten terms since many terms are not shared between the smoothing methods.

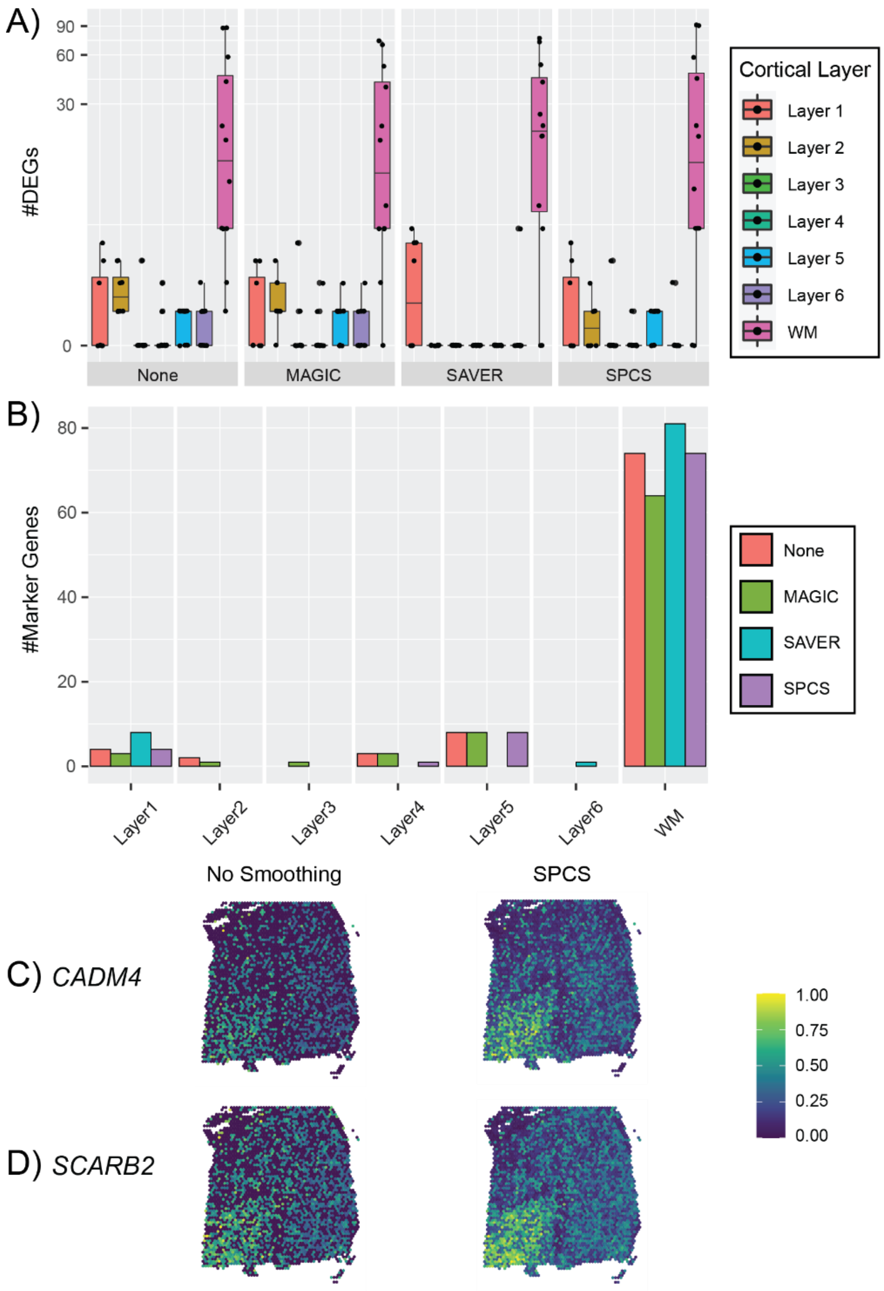


**Supplementary Figure 4.** Biological analysis of DLPFC slides with no smoothing, MAGIC, SAVER, and SPCS smoothing methods. (a) Number of DEGs identified in each slide and cortical layer. (b) The number of shared significant DEGs between each smoothing method and a list of cortical layer marker genes [11]. (c-d) Heatmap of unsmoothed and SPCS smoothed expression of (c) *CADM4* and (d) *SCARB2* in sample 151673. For demonstration purposes, expression of genes was linear transformed into range of 0 to 1 as normalization.
